# Convergent evolution of sensory palps in Scalibregmatidae (Annelida)

**DOI:** 10.1101/2025.10.09.681389

**Authors:** Paul Kalke, Katrine Worsaae, Patrick Beckers, Alejandro Martinez Garcia, Conrad Helm

**Affiliations:** Helm Lab, Johann-Friedrich-Blumenbach-Institute, Animal Evolution and Biodiversity, University of Göttingen, Göttingen, Germany; Marine Biological Section, Department of Biology, University of Copenhagen, Copenhagen, Denmark; Institute of Evolutionary Biology and Zooecology, Evolutionary Biology and Ecology, University of Bonn, Bonn, Germany; CNR - IRSA: Water Research Institute, Verbania, Italy

**Keywords:** head appendages, convergent evolution, Scalibregmatidae, nervous system, musculature, cLSM

## Abstract

Scalibregmatidae are deposit feeders inhabiting muddy or sandy sea floors at considerable depths, and usually lack head appendages common in many other marine annelids. Surprisingly, one lineage within this family bears palp-like structures comparable to anterior head appendages in other Annelida. Nevertheless, a homology of the scalibregmatid appendages with those of other annelids is highly questionable. Using an integrative morphological approach including immunohistochemistry, azan-histology, and 3D reconstruction, we examined the neural innervation of the anterior head region in one travisiid and two scalibregmatid species: *Travisia* sp. (without appendages), *Scalibregma celticum* (lateral, prostomial horns), and *Axiokebuita cavernicola* (palp-like appendages). *Scalibregma and Axiokebuita* show an innervation pattern of their appendages different from that of palps in other annelids, while no prominent prostomial innervation is present in *Travisia*. The short, lateral horn-like appendages of *S. celticum* possess a quite diffuse neuronal scaffold extending from distinct neurite bundles originating from multiple regions of the brain. The elongated palp-like head appendages of *A. cavernicola* exhibit a neuronal innervation pattern comparable with that of thin sensory antennae known from other annelid groups. Thus, innervating neurites solely originate in the dorsal part of the brain, a pattern not observable for “true” feeding- or sensory palps of other taxa. We also shed new light on the mysterious “neck organ” in scalibregmatids, which are shown to be ventral dublications of the olfactory nuchal organs considering the level of external features and nervous innervation. These results, combined to previous phylogenetic studies, show that Scalibregmatidae are the only so far known annelids to have re-evolved palp-like sensory head structures after a secondary loss. Thus, this is the first example of convergent evolution of palps, a character complex hitherto showing similar overall innervation patterns and considered homologous across all annelids.

## 1. Introduction

The globally distributed annelid family Scalibregmatidae comprises around 141 species in 15 genera (Blake 2020; Blake 2025). Most species are subsurface deposit feeders inhabiting muddy or sandy sediments, typically at considerable depths or high latitudes (Parapar et al. 2011; Blake 2020; Blake 2025). While a burrowing lifestyle is typical for the family, the *Axiokebuita–Speleobregma* clade represents a striking ecological shift, inhabiting crevicular and gravelly habitats in caves and the deep sea (Martínez et al. 2013). These environments, defined by high permeability and inertial water flow, directly correlates with their exceptional morphology (Martínez et al. 2014), and contrast with the low-permeability and porous water flow dominating muddy and sandy substrates.

Despite decades of taxonomic revisions and the reallocation of taxa (Blake & Kudenov 1981; Kudenov & Blake 1985; Pleijel & Fauchald 1993; Paul et al. 2010; Blake & Maciolek 2020), the phylogenetic position of Scalibregmatidae and the relationships among its genera remain only partly resolved. Traditionally classified within Scolecida, sister group to the remaining annelids, the family is now consistently recovered as highly derived within the Sedentaria clade, forming a group with Travisiidae, Maldanidae and Arenicolidae (Rouse & Fauchald 1997; Rouse & Pleijel 2001; Bleidorn et al. 2003, 2005; Persson & Pleijel 2005; Struck et al. 2007; Paul et al. 2010; Martínez et al. 2014; Martín-Durán et al. 2021). Within the family, *Axiokebuita* and *Speleobregma* constitute a derived clade (see Figure 1).

**Figrue 1:**
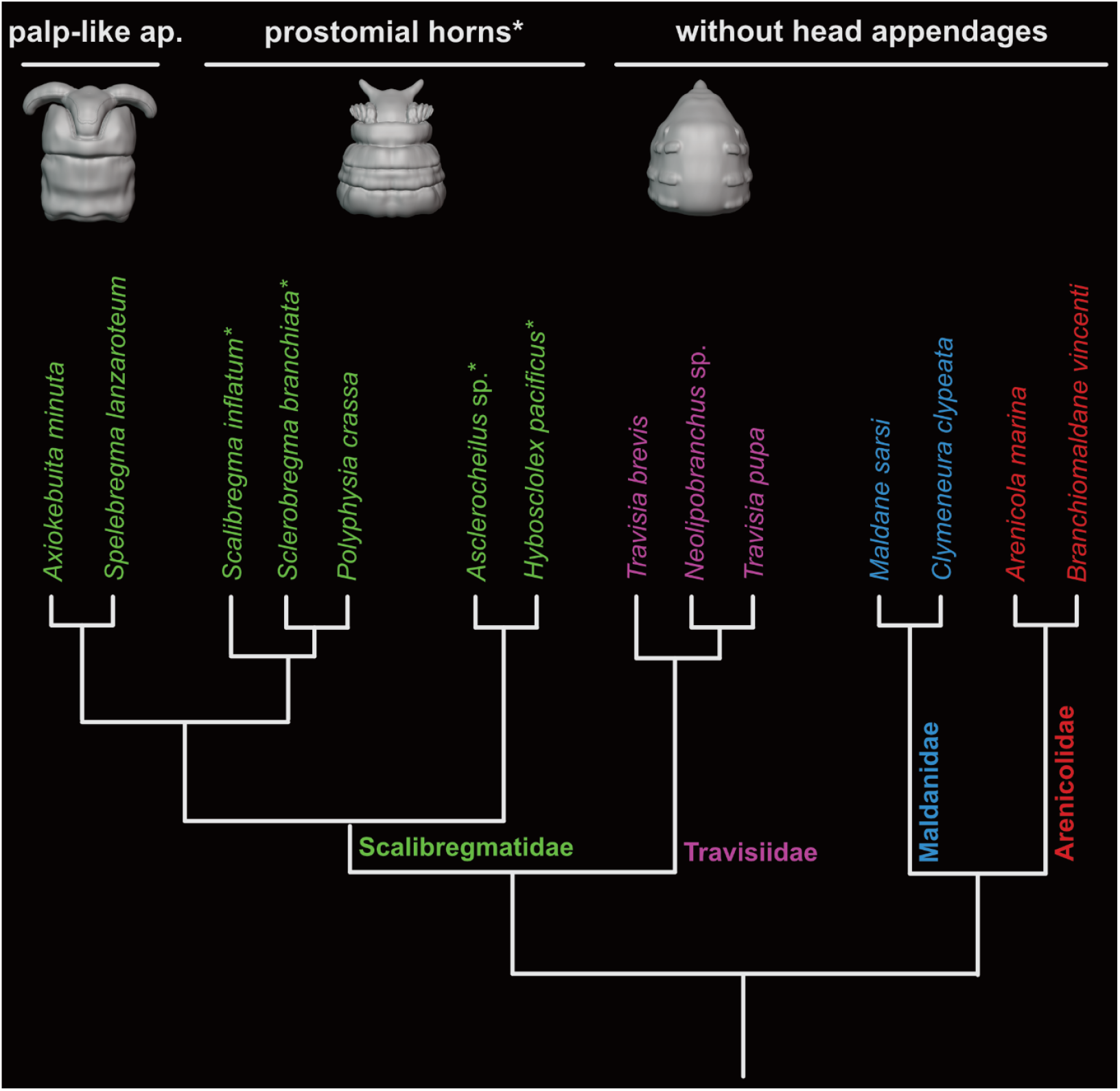
Phylogenetic relationship of Scalibregmatidae, Travisiidae, Maldanidae and Arenicolidae, modified after Martínez et al. (2014). The external morphology of anterior ends of investigated *Travisia* sp., *Scalibregma celticum* and *Axiokebuita cavernicola* are shown exemplarily for their group as 3D - models. Members of Scalibregmatidae are highlighted in green, Travisiidae in pink, Maldanidae in cyan and Arenicolidae in red. Scalibregmatid species processing prostomial/lateral horns are marked with asterisks.

Morphologically, Scalibregmatidae are characterized by a rugose epidermis, segmental annuli, and a T-shaped prostomium with presumably sensory lateral horns, which can be absent or modified in several taxa (Rouse & Pleijel 2001; Persson & Pleijel 2005). In *Axiokebuita* and *Speleobregma* the prostomial appendages are elongated and broad and positioned ventrally. These, so-called palps, are densely ciliated and produce water currents, putatively used to gather food particles (Bertelsen 1986; Pocklington & Fournier 1987; Persson & Pleijel 2005). *Axiokebuita* and *Speleobregma* are further characterized by presence of an adhesive pygidium, and absence of lyrate chaetae and branchiae that characterize other scalibregmatids (Martínez et al. 2013). Recent work has revealed additional overlooked scalibregmatid structures, including epidermal ornamentation, ciliary patterns, and glandular arrangements, leading to refined species diagnoses (Martínez et al. 2013; Blake 2015). Particularly enigmatic is the so-called “neck organ,” described in *Axiokebuita* as a paired ciliated peristomial structure of potential sensory function (Persson & Pleijel 2005). Its evolutionary significance remains unresolved, making it a warranting structure for deeper investigation.

Multiple studies have investigated and/or debated the homology of palps and other head appendages across annelids (Orrhage 1980, 1991, 1995, 1996; Rouse & Fauchald 1997; Orrhage & Müller 2005; Martinez et al. 2014; Beckers et al. 2019a, b; Helm et al. 2022; Kalke et al. 2021, 2024). Recent studies of representatives from all major clades on the annelid tree of life (after Weigert et al. 2014), such as Paleoannelida (Beckers at al. 2019a, b; Kalke et al. 2024), Chaetopteriformia (Helm et al. 2022), Amphinomidae (Beckers & Tilic, 2021; Kalke et al. 2024), Errantia such as Syllidae (Schmidbaur et al. 2020) and Eunicida (Kuhl et al. 2022), and Sedentaria including Siboglinidae (Worsaae et al. 2016; Rimskaya-Korsakova et al. 2018), Spionidae (Kalke et al. 2024) and Terebelliformia (Kalke et al. 2021), have supported that all the structures described as palps so far share the same innervation pattern. Neuroanatomical studies on Scalibregmatidae are scarce, to date, only Orrhage & Müller (2005) have reported that a palp innervation is present even in scalibregmatid taxa lacking protruding palps or prostomial horns. An ancestral character estimation on the scalibregmatids’ molecular tree suggests that ventral, ciliated and ungrooved palps in *Axiokebuita–Speleobregma* evolved from ancestors with prostomial horns (Martínez et al. 2014), although their homology have not been anatomically investigated. Based on those results, and lacking further morphological evidence, the scalibregmatid appendages can moreover either be interpreted as a reversal to the plesiomorphic annelid state ‘prostomial palps present’, or as a convergent gain after secondary entire loss of palps in the burrowing ancestor of Maldanidae, Arenicolidae, and Travisiidae, as suggested by other authors (Rouse & Fauchald 1997; Struck 2011).

These doubts on homology of scalibregmatid prostomial appendages underline the need for more anatomical data and a detailed re-evaluation of the evolutionary history of palps in annelids. Especially the derived position of Scalibregmatidae within Sedentaria (Martín-Durán et al. 2021), as sister group to burrowing direct deposit feeding taxa like Travisiidae, Maldanidae, and Arenicolidae, without head appendages, make them an ideal model to investigate a potential reversal to or convergent formation of palp-like appendages. Hence, our study provides a detailed investigation of the nervous system and anatomy of the anterior region of *Travisia* sp., *Scalibregma celticum* (with prostomial horns) and *Axiokebuita cavernicola* (with palp-like appendages). We use an integrative morphological approach including immunostaining and confocal laser scanning microscopy (CLSM) as well as azan-stained histological sections with subsequent 3D-visualisation. With this comparative approach, we will analyze the evolutionary history of the head appendages of these enigmatic annelids and test the homology or potential first convergent evolution of palp-like head appendages in annelids. We moreover discuss functional aspects of prostomial horns and palp-like appendages in the categories of feeding vs sensory. In addition, we tackle the question about the evolution and function of the enigmatic “neck organ” and further newly identified ciliary structures.

## 2. Material and Methods

### 2.1 Specimen collection

Adult specimens of *Travisia sp*. (Johnston, 1840) were collected in 2013 in Ferrol (Spain) via a bottom dredge. Adult *Scalibregma celticum* (Mackie, 1991) were collected via a bottom sampling dredge at around 20 m depth in the Bay de Morlaix near the biological field station (Roscoff, France) in 2020. Adult specimens of *Axiokebuita cavernicola* (Martínez, Di Domenico & Worsaae, 2013) were collected by scuba diving from the Cerebros Cave, Tenerife, Canary Islands at around 8m depth. For more detailed information on the locality check Martinez et al. (2013).

### 2.2 Azan staining, histological sectioning and 3D-reconstruction

Semi-thin sections and AZAN staining of adult *Travisia* sp., *Scalibregma celticum*, and *Axiokebuita cavernicola* were prepared following the protocol of Beckers et al. (2013). Specimens were first relaxed in 7% MgCl_2_ and subsequently fixed in Bouin’s fluid for 12 h. They were then dehydrated through an ethanol series and transferred into methylbenzoate and butanol. Pre-incubation was carried out in Histoplast (Thermo Scientific, Dreieich, Germany), and embedding was done in Paraplast (McCormick Scientific, Richmond, USA). Serial sections of 5 μm thickness were cut using a Reichert-Jung Autocut 2050 microtome (Leica, Wetzlar, Germany) and mounted on albumen–glycerin coated glass slides. Sections were stained with Carmalaun, differentiated in 5% sodium phosphotungstate, rinsed in distilled water, counterstained with aniline blue–orange G, and finally embedded in Malinol (Waldeck, Münster, Germany). Under Azan staining, the neuropil of the nervous system appears gray, cell nuclei red, the extracellular matrix blue, and musculature orange (Beckers et al., 2013). Digitalization of sections was performed at 40× magnification using a slide scanner [Olympus dotSlide 2.2, Olympus, Hamburg], followed by alignment with IMOD (Kremer et al., 1996) and imodalign. For 3D reconstruction, a digital workflow exclusively based on open-source software was applied, including ImageJ, MeshLab, and Blender (Kalke & Helm, 2022). For schematic 3D renderings, curve tools in Blender were used (see tutorial: https://www.youtube.com/watch?v=Ve9h7-E8EuM

### 2.3 Immunohistochemistry

Anatomical details of the scalibregmatids *Scalibregma celticum* and *Axiokebuita cavernicola* were investigated using standard immunohistochemical staining protocols. Hence, a minimum of 5 specimens of each species were relaxed in 7% MgCl2 and subsequently fixed in 4% paraformaldehyde (PFA) in 1x phosphate buffered saline with Tween (PTW = 1x PBS: 0.05 M PBS / 0.3 M NaCl / 0.3% TritonX. Fixation was performed at room temperature (RT) for 2 h. After fixing, the specimens were washed and stored in PTW containing 0,005% NaN3 until usage at 4°C. For antibody staining, specimens were rinsed 3 × 10 min in PTW and incubated in Collagenase D (from Clostridium histolyticum, Roche Diagnostics GmbH, Mannheim, Germany, 0,24U/mg lyo.) and 10 μg proteinase K/ml PTW. Adult specimens were incubated in Collagenase D solution (10mg/ml) in PTW + 0,3% Triton X-100 for 1 hour at RT, rinsed minimum 3 times for 5 min in PTW and subsequently incubated in proteinase-K for 20 min. To stop the proteinase-K reaction, samples were rinsed twice in glycine (2 mg glycine/ml PTW), and washed 3 × 5min in PTW. Afterwards, the specimens were re-fixed using 4% PFA in PTW containing 0.6/0.3% TritonX for 20 min at RT. Subsequently, the samples were rinsed 2 × 5min in PTW as well as 2 × 5min in THT (0.1 M TrisCl, 0.3 TritonX, pH 8,5), and blocked with 5% goat serum (Sigma-Aldrich, Steinheim, 25 μL goat serum in 500 μL THT) for 2 h. For the visualization of the nervous system specimens were incubated with the primary antibodies against α-tubulin (Anti-acetyl α - tubulin, clone 6-11B-1, Merck, Darmstadt, 2 μL tubulin in 500 μL incl. 5% goat serum) and serotonin (5-HT-serotonin), ImmunoStar Inc., Husk, USA, 1 μL in 500 μL incl. 5% goat serum) in THT for 48–72 h at 4°C. Afterwards, samples were rinsed 2 × 10 min in 1 M NaCl (to deactivate the primary antibody) in THT and washed 5 × 30 min in THT. Subsequently, the samples were incubated in the secondary antibodies goat-anti-mouse 633 (Alexa Fluor® 633 goat-anti-mouse IgG (H + L), Thermo Fisher Scientific Inc., Waltham, USA, 1 μL in 500 μL inlc. 5% goat serum) and goatanti-rabbit 488 (Alexa Fluor® 488 goat-anti-rabbit IgG (H + L), Thermo Fisher Scientific Inc., Waltham, USA, 1 μL in 500 μL incl. 5% goat serum) in THT for 48–72 h at 4°C. After staining, specimens were rinsed 5 × 30 min in THT and 2 × 5 min in PTW. For the visualization of the musculature specimens were incubated in rhodamine-labeled phalloidin (Thermo Fisher Scientific; 5 µl methanolic stock solution in 500 µl inlc. 5% goat serum) in THT overnight at 4°C. Additionally, all samples were incubated in DAPI (DAPI, Thermo Fisher Scientific Inc., Waltham, USA, 5 μL in PTW) in PTW (we observed better results for DAPI-stainings in PTW) overnight at 4°C. Subsequently, the specimens were dehydrated in an ascending isopropanol series, cleared using Murray’s clear (benzyl alcohol & benzyl benzoate, 1:2) and embedded between two cover slips using DPX mounting medium (Merck, Darmstadt, Germany). The specimens were analyzed with the confocal laser scanning microscopes Leica TCS SP8 (Leica Microsystems, Wetzlar, Germany). The confocal image stacks were processed with Leica AS AF v2.3.5 (Leica Microsystems) and Imaris × 64 9.5.0 (Bitplane AG, Zurich, Switzerland).

## 3. Results

The anterior adult morphology with focus on nervous system and musculature of *Travisia sp*. is shown in Figure 2, of *Scalibregma celticum* in Figure 3 and adult *Axiokebuita cavernicola*. in Figure 4. A overall summary of the data including a simplified phylogenetic tree (modified after Martínez et al. 2014) and 3D-visualisations, with related videos as supplementary files, is provided in Figure 5.

**Figure 2.**
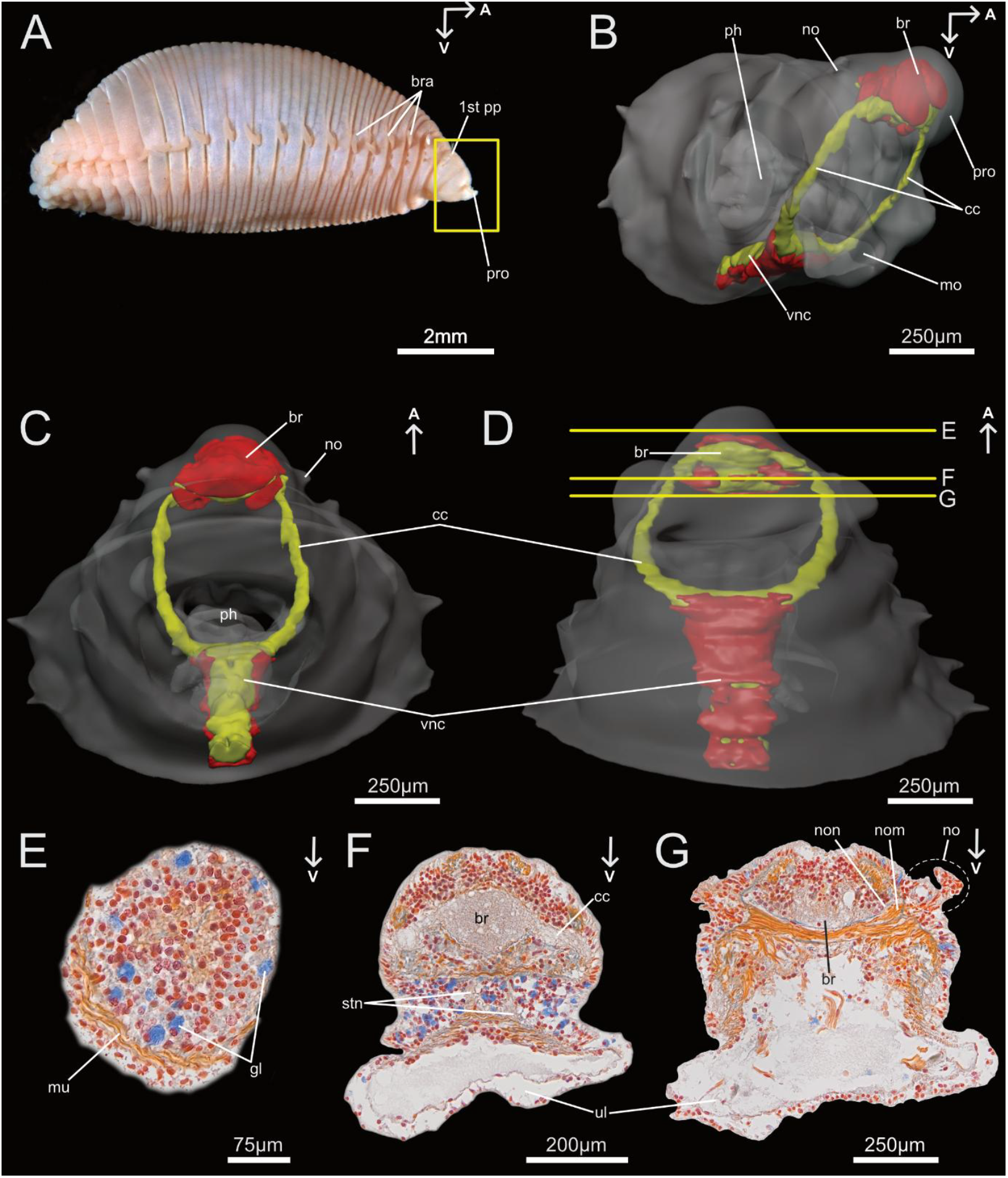
Anterior Morphology of *Travisia sp*. (Travisiidae). A) Light microscopic image of *Travisia forbesii* (Johnston, 1840) in lateral view, shown is the classic maggot -l ike habitus of the genus *Travisia*. B) 3D-reconstruction of the anterior nervous system of *Travisia*, in antero-lateral view, with the condensed brain (*br*), circumoesophageal connectives (cc) and ventral nerve cord (*vnc*). C) Same reconstruction in ventral view, see the perikarya of the brain are located dorsally to it s neuropil, the one of the *vnc* are located ventrally to it, and the pointy bumps of the nuchal organs (*no*). D) 3 D-reconstruction of *Travisia* in ventral view, the yellow transversal lines mark the position of the histological cross-section shown in 2E, F and G. E) Azan-stained cross-section trough the pointy prostomium without any sign of nervous system structures. F) Cross-section through the condensed brain (*br*) show its transition to the the *cc*, stomatogastric neurite bundles can be distinguished. G) Cross-section through the posterior part of the *br* with the nuchal organ nerve (*non*), the nuchal organ retractor muscle (*nom*) and the bump-like *no*. 1st pp - first parapodium, br – brain, bra – branchiae, cc – circumoesophageal connectives, gl – glandular cells, mo – mouth opening, mu – musculature, no – nuchal organ, non – nuchal organ nerve, ph – pharynx, pro – prostomium, ul – upper lip, vnc – ventral nerve cord

**Figure 3.**
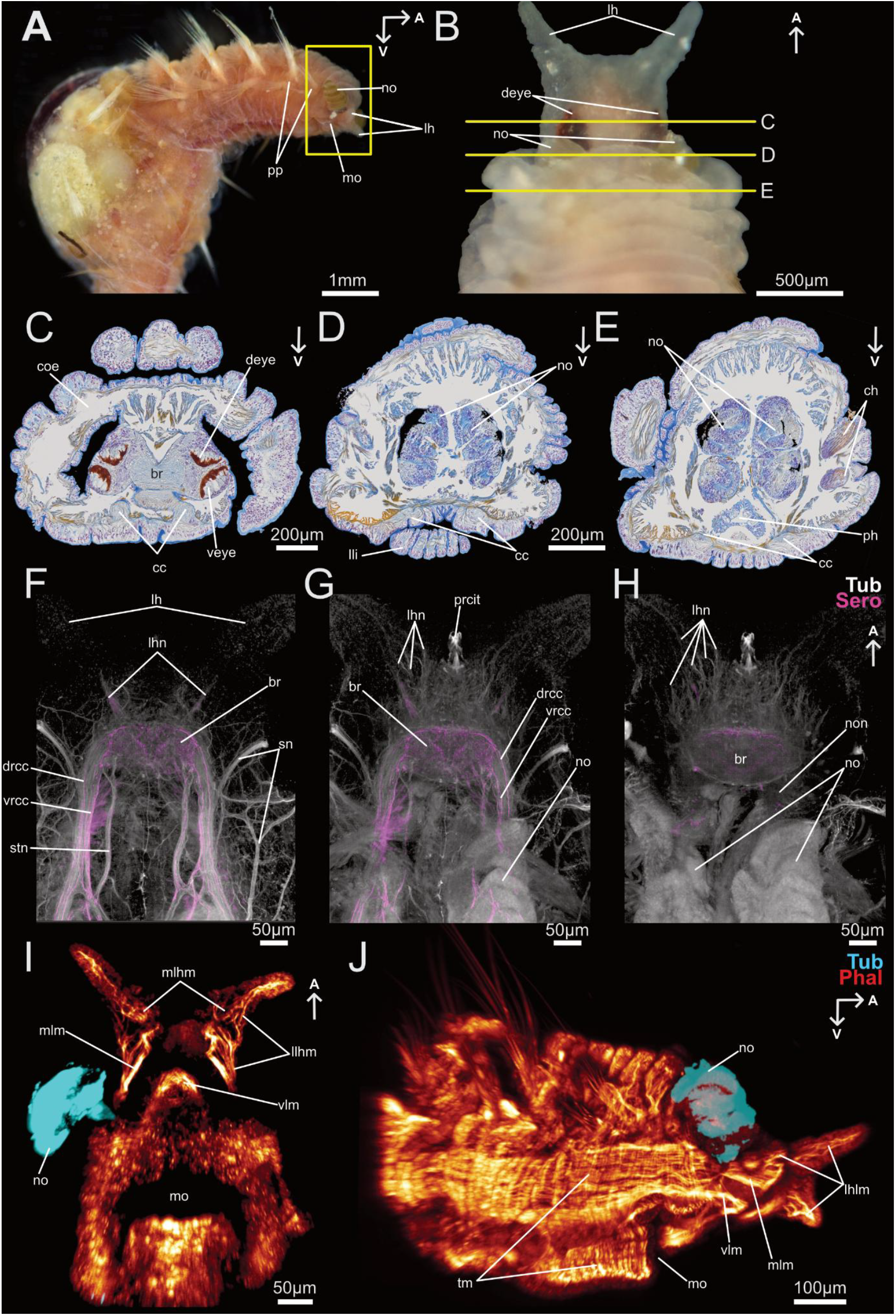
Anterior morphology of *Scalibregma celticum*. A) Light microscopic image (©Patrick Beckers) of *S. celticum* in ventro-lateral view, the curly nuchal organs (*no*) are everted and the prostomial prostomial horns can be distinguished. B) Light microscopic image (©Patrick Beckers) of the prostomium with prostomial horns and peristomium in dorsal view with inverted *no* and prominent, pigmented dorsal eyes. The yellow transversal lines mark the position of the Azan-stained histological cross-sections 3C, D and E. C) Histological cross-section through the transition area of pro - to peristomium with the posterior brain region (*br*), the circumoesophageal connectives (*cc*) and the pairs of ventral (*veye*) and dorsal eyes (*deye*). D) Histological cross-section through the inverted *no* sitting in pouches. E) Histological cross-section through the posterior end of the nuchal organ pouches, chaetae of neuro - and notopodium of the first parapodia are shown. F, G, H) Digital cross-section of confocal images with α-tubulinergic (white) and serotonergic (pink) immunoreactivity, through the prostomium, in ventro - dorsal progression. F) Serotonergic signal of the two roots (*vrcc* and *drcc*) of the *cc* as well as the ventral brain (*br*) region and the two serotonergic nerve of the *lh* are shown. In dorsal progression (G and H), the serotonergic signal in the brain neuropil decreases and the number of prostomial horn nerves (*lhn*) from the dorsal brain region increase. In the center of the prostomium a ciliary tuft with i ts innervation by the dorsal brain region can be distinguished. I, J) Confocal images of the anterior phalloidin signal, in ventral view in I and lateral view in J. I) The musculature inserting in the prostomial horns can be distinguished (*mlhm* and *lhlm*). J) Details l ike insertion of the *mlm* and *tm* are shown. br – brain, cc – circumoesophageal connectives, ch – chaetae, coe – coelom, deye – dorsal eyes, drcc – dorsal root of cc, lh – prostomial horn, lhlm – prostomial horn longitudinal muscle, lhn – prostomial horn nerves, ll i – lower l ip, mlhm – median prostomial horn muscle, mlm – median longitudinal muscle, mo – mouth opening, no – nuchal organ, non – nuchal organ nerve, ph – pharynx, pp – parapodia, prcit – prostomial ciliary tuft, sn – segmental neurite bundle, stn – stomatogastric neurite bundles, tm – transversal muscle fibers, veye – ventral eyes, vlm – ventral longitudinal muscle, vnc – ventral nerve cord, vrcc – ventral root of cc

**Figure 4.**
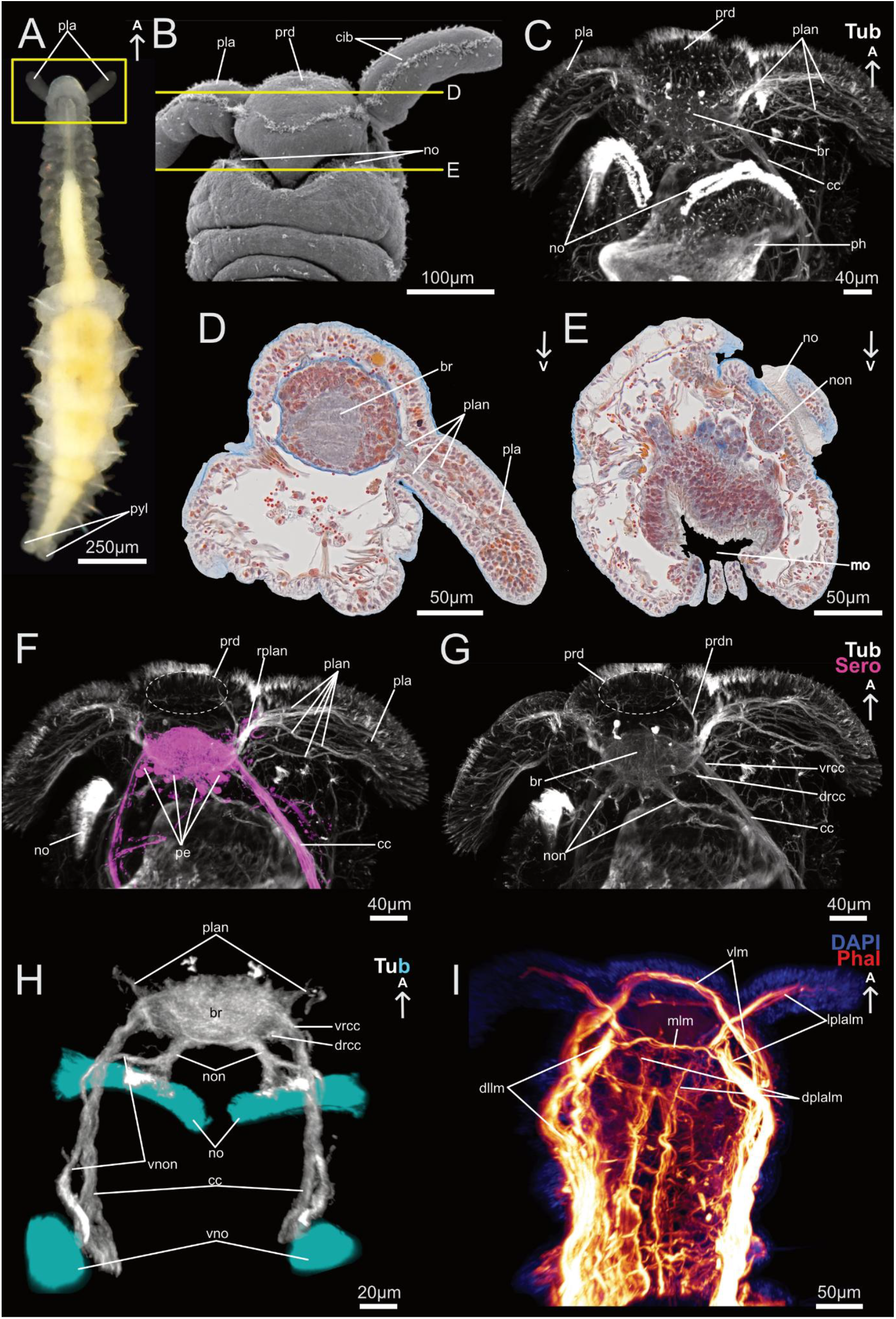
Anterior morphology of *Axiokebuita cavernicola*. A) Light microscopic image in dorsal view of small specimen with regenerating anterior end – by Arne Nygren. B) SEM image in dorsal view with ciliary structures, palp-like appendages (*pla*) and prostomial depression (*prd*), modified after Parapar at al. (2021). C) Confocal image of α-tubulinergic anterior nervous system in dorsal view with prominent nuchal organs (*no*). D) Histological Azan-stained cross-section through one *pla* with i ts nerves (*plan*) leaving the brain (*br*). E) Histological cross-section through the huge *no*, ciliary pit and nuchal organ nerve (*non*) can be distinguished. F) confocal image of serotonergic and α-tubulinergic anterior nervous system with common origin of *plan* from the dorsal brain region (*rplan* and huge serotonergic perikarya (*pe*). G) Same confocal image (only α-tubulin) with highlighted *prd* and its innervation (*prdn*). H) Confocal image in dorsal view with *no* and ventral nuchal organ (*vno*) i ts innervation (*non, vnon*). I) Confocal image of the phalloidin signal of the anterior musculature in ventral view, with two muscle inserting into the *pla* (*dplalm, lplalm*). br – brain, cc – circumoesophageal connectives, cib – ciliary bands, dllm – dorso-lateral longitudinal muscle, dplalm – dorsal palp-l ike appendage longitudinal muscle, drcc – dorsal root of cc, lplalm – lateral palp-like appendage longitudinal muscle, mlm – median longitudinal muscle, mo – mouth opening, no – nuchal organ, non – nuchal organ nerve, pe – serotonergic perikarya, ph – pharynx, pla – palp-like appendage, plan – palp-like appendages nerves, prd – prostomial depression, prdn – prostomial depression nerves, pyl – pygidial lobes, rplan – root of palp-like appendages nerves, tm – transversal muscle f ibers, vlm – ventral longitudinal muscle, vnc – ventral nerve cord, vno – ventral nuchal organ, vnon – ventral nuchal organ nerve, vrcc – ventral root of cc

**Figure 5.**
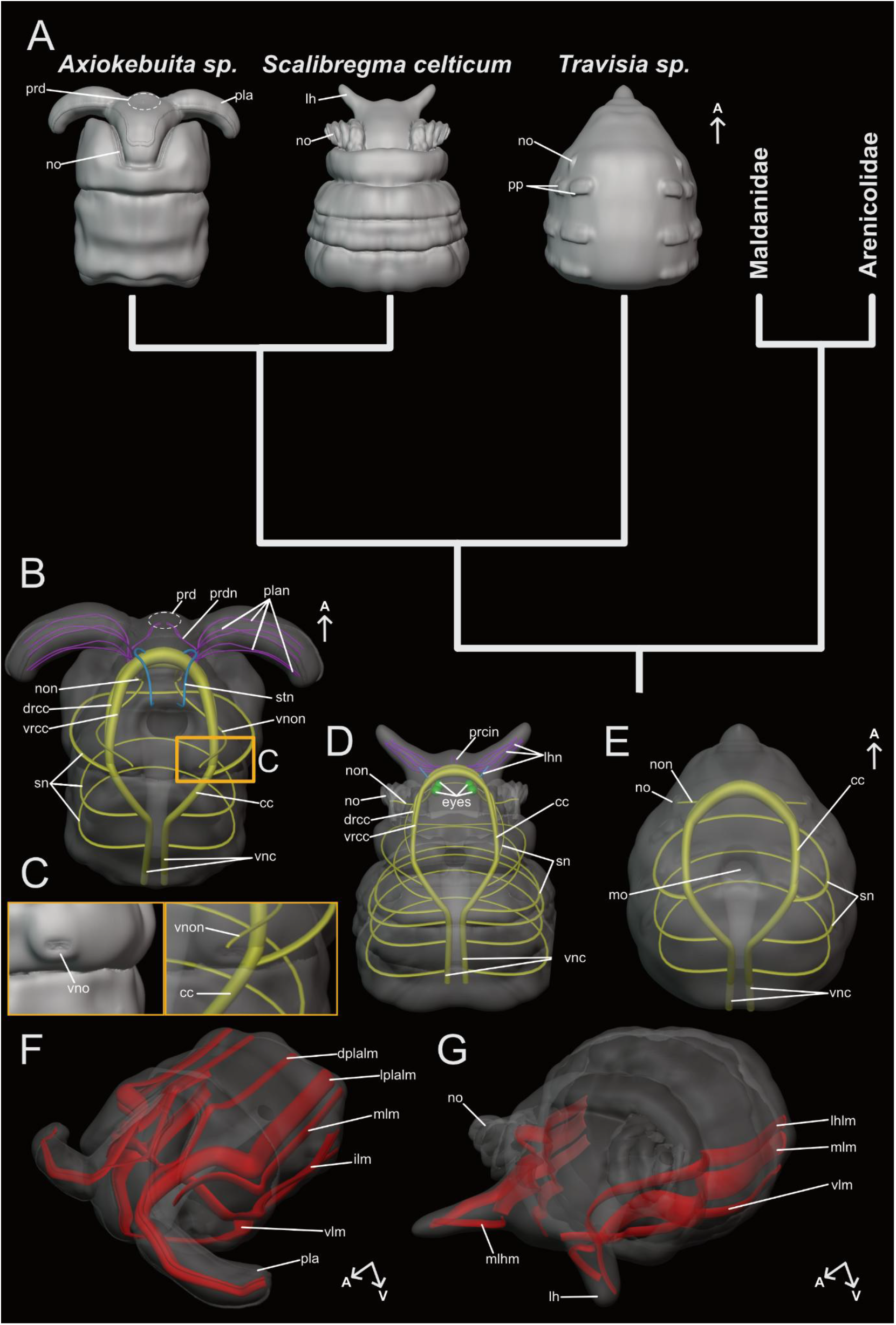
Summary of our results concerning the anterior morphology of *Travisia sp*., *Scalibregma celticum* and *Axiokebuita cavernicola*. A) Simplified Phylogeny modified after Martínez et al. (2014) with prominent external features in 3 d-models in dorsal view. B) Summery of the anterior nervous system in a 3D-model of *A. cavernicola*, in ventral view. Neurite bundles originate solely from th e *vrcc* are highlighted in cyan, from the *drcc* in violet and nuchal organ nerves in yellow. C) External morphology and innervation of the ventral nuchal organ of *A. cavernicola*. D) Summary of the anterior nervous system in a 3D-model of *S. celticum*, in ventral view, number of prostomial/ lateral horn nerves (*lhn*) is strongly reduced. Neurite bundles originate solely from the *vrcc* are highlighted in cyan, from the *drcc* in violet and nuchal organ nerves in yellow. E) Summary of the anterior nervous system in a 3D-model of *Travisia sp*., in ventral view. F) Summary of the anterior musculature of *A. cavernicola*, transversal musculature omitted. G) Summary of the anterior musculature of *S. celticum*, transversal musculature omitted.

For general terms and annotations of the nervous system characters, we refer to Richter et al. (2010), and Kalke et al. (2024). For the muscle characters we refer to Weidhase et al. (2016). Annotations for external features were used after Martinez et al. (2021) and Parapar et al. (2011).

### 3.1 The anterior nervous system of *Travisia sp*. (Travisiidae)

Members of the genus *Travisia* have a maggot-like body (here *Travisia forbesii*, Figure 2A) small anterior parapodia (*pp*) and a pointed prostomium (*pro*) without appendages or conspicuous ciliation. Prominent eye spots are absent. The anterior nervous system consists of a condensed brain neuropil (Figure 2B, C, D) and its perikarya are mostly arranged dorsal of the neuropil. In the tip of the pointy prostomium prominent musculature (*mu*) and gland-like cells (*gl*) are present, but no neurite bundles or other neuronal structures extend to the anterior end (Figure 2E). The brain neuropil (*br*) appears condensed and unstructured, a dorsal or ventral root of the circumoesophageal connectives (*cc*), as well as distinct tracts, cannot be distinguished in the histological sections (Figure 2F). The same condensed architecture can be found in the *cc*, which surround the pharynx (*ph*) and merge into the subsequent ventral nerve cord (*vnc*), where the perikarya are located ventrally to the neuropil (Figure 2B, C and D). The minute nuchal organs (*no*), externally visible as a pointy swelling, lie dorso-lateral to the brain neuropil at the prostomium peristomium boundary (Figure 2G, see also 5E). Although prominent ciliation is absent, each organ presents characteristic retractor muscle (*nom*) and is innervated by the nuchal organ nerve (*non*) originating posterior of the brain neuropil (Figure 2G, see also 5E).

### 3.2 The anterior nervous system and musculature of *Scalibregma celticum* (Scalibregmatidae)

*Scalibregma celticum* is a pinkish annelid with prominent eye spots (*deye, veye*) (Figure 3B, C), curled nuchal organs (*no*) that can retract into peristomial pouches (Figure 3A, B, D, E, G, H, I and J) and a T-shaped prostomium with two prominent prostomial, lateral horns (Figure 3A, B and F). The four cerebral eye spots (*veye, deye*) are rather large, pigmented and intraepidermal, located at the boundary of the first broad peristomial collar (Figure 3B, C, and 5D). Posteriorly, the nuchal organs and their peristomial pouches connect to the brain via the nuchal-organ nerve (*non*) (Figure 3H, 5D). The anterior central nervous system is complex and very condensed, with a dorsal protrusion of the brain (*br*) neuropil (Figure 3F-H, 5D). Distinguishing the ventral (*vrcc*) and dorsal roots (*drcc*) of the circumoesophageal connectives (*cc*) is only possible using serotonin-like immunoreactivity (Figure 3F-H). In the ventral-most digital section through the brain and *cc* region (Figure 3F), serotonergic stomatogastric neurite bundles extend posteriorly and a single serotonergic prostomial-horn nerve (*lhn*) runs anteriorly (Figure 3F, 5D). Slightly more dorsally (Figure 3G), a prostomial ciliary tuft (*prcit*) and its innervation (see also Figure 5D) from the condensed dorsal brain region, together with several prostomial horn nerves are visible. In the dorsal-most digital section, aligned with the *no* and their nerve *non* (Figure 3H) an increasing number of *lhn* project into the prostomial horns; these originate broadly from the dorsal brain region in an unorganized manner (see also Figure 5D).

The anterior musculature is dominated by laterally-position longitudinal muscle fibers (Figure 3I, J and 5G), whereas very slim, regularly arranged transverse muscles encircle the body (Figure 3I, J). The ventral longitudinal muscles (*vlm*) surround the mouth opening and, forming a semicircle, insert ventrally to the prominent median longitudinal muscle (*mlm*). The *mlm* inserts the prostomium directly ventral to the prostomial horns. The third, most dorsal bundle is the prostomial horn longitudinal muscle (*lhlm*), which inserts at the lateral tip of each prostomial horns (Figure 3I, J and 5G). A second bundle inserts at the medial tip of the prostomial horn (*mlhm*) and terminates at its base, close to the *mlm* insertion (Figure 3I and 5G).

### 3.3 The anterior nervous system and musculature of *Axiokebuita cavernicola* (Scalibregmatidae)

*Axiokebuita cavernicola* has an elongated body with two prominent adhesive phygidial lobes and two palp-like appendages (*pla*) at the anterior end, pigmented eye spots are missing (Figure 4A, B). Two transversal ciliary bands (*cib*) proceed from one tip of the palp-like appendages to the other by crossing the prostomium (Figure 4B, 5A). At the center of the prostomium a prostomial depression (*prd*) can be distinguished, while a deep furca, marks the transition to the peristomium and in which the prominent nuchal organ is located (Figure 4B, C, E and H, 5A). Like in *Travisia sp*. and *S. celticum* the brain is condensed, and the ventral (*vrcc*) and dorsal (*drcc*) root of the circumoesophageal connectives (*cc*) are only detectable along the *cc*, but not within the brain neuropil (Figure 4C, D, F-H, 5B). The perikaya are located dorsally and posterior to the neuropil including huge serotonergic perikarya (4D, F). The *prd* is innervated be nerves originating from the dorsal brain region (*prdn*), proceeding parallel to the palp-like appendage nerve root (*rplan*) and split up anteriorly (Figure 4B, C, F and G, 5B). Additionally, all five palp-like appendage nerves (*plan*) originate from the dorsal brain region (solely from one root) and start to split up in antero-lateral direction, innervating the whole *pla* (Figure 4C, D, F-H, 5B).

The prominent nuchal organs are innervated by two huge nerves (*non*) originating from the posterior margin of the dorsal brain region (Figure 4E, G, H and 5B). Before entering the nuchal organ these nerves branch to give rise to the ventral nuchal organ nerve (*vnon*), which proceeds in ventro-lateral direction and terminates in two ciliated pits posterior to the mouth opening - the ventral nuchal organs (*vno*) or so-called “neck organ” (Figure 4H and 5B, C).

The anterior musculature of *A. cavernicola* shows a comparable pattern as observable in *S. celticum*. Thus, its dominated by longitudinal muscle fibers, being accompanied by sparse, very thin transverse fibers (*tm*) (Figure 4J). A ventral longitudinal muscle (*vlm*) runs along the ventro-lateral side of the body and inserts into the prostomium anteriorly of the mouth opening (Figure 4I, 5F). Dorsal to this, the intermediate longitudinal muscle (ilm) encircle the mouth opening and terminates dorsal to it. The median longitudinal muscle (*mlm*) runs parallel to the other muscle and inserts into the prostomium at the base of the palp-like appendages. Dorsally to it, and parallel, the lateral palp-like longitudinal muscle appears (*lplalm*). The dorsal-most pair of muscle bundles is represented by the dorsal palp-like appendage longitudinal muscle (*dplalm*), which joins the progression of *lplalm* to the tip of the *pla*.

## 4. Discussion

Palps play a crucial role in annelid evolution, yet their evolutionary history remains controversial (Rouse & Fauchald 1997; Orrhage & Müller 2005; Martínez et al. 2014; Beckers et al. 2019a, b; Schmidbaur et al. 2020; Helm et al. 2022; Kalke et al. 2021, 2024). Recent studies emphasize that external features alone are insufficient for defining head appendages, and that their neuronal as well as muscular innervation patterns must also be considered. For example, in some eunicids, palps and antennae are morphologically similar but can be distinguished by their distinct innervation pattern (Kuhl et al. 2022). Generally, palps in annelids are always innervated by nerves originating from both the dorsal and ventral roots of the circumoesophageal connectives (*drcc, vrcc*), whereas antennae show neuronal innervation only by neurites originating from the dorsal root (Beckers et al. 2019a, b; Schmidbaur et al. 2020; Kuhl et al. 2022; Kalke et al. 2021, 2024).

Although the precise role of short sensorial palps in errant taxa remains debated and comparative datasets are still quite scarce, anatomical investigations across Paleoannelida (Beckers et al. 2019a, b), Chaetopteriformia (Helm et al. 2022), Amphinomidae (Beckers & Tilic 2022; Kalke et al. 2024) and several members of the Sedentaria (Worsaae et al. 2016; Rimskaya-Korsakova et al. 2018; Kalke et al. 2021, 2024) – support the hypothesis that, at least all groups with feeding-palps examined so far conctitute a homologous character complex within Annelida.

In Scalibregmatidae, the interpretation of prostomial/lateral horns and palp-like appendages is closely linked to existing phylogenetic hypotheses and associated to ecological transitions (Martínez et al. 2013, 2014). Martínez et al. (2014) outlined two scenarios: a reversal of the plesiomorphic condition under the Palpata–Scolecida hypothesis, or a regain after secondary loss in the burrowing ancestor of Maldanidae, Arenicolidae, and Travisiidae, consistent with the Errantia–Sedentaria hypothesis. We favor the latter, which is supported by several independent molecular studies (Struck et al. 2011; Weigert et al. 2014; Laumer et al. 2015; Struck et al. 2015; Weigert & Bleidorn 2016), in which repeated secondary losses of palps are inferred in Orbiniidae (Helm et al. 2015), Opheliidae (Parapar et al. 2021), Dinophilidae (Windoffer & Westheide 1988; Kerbl et al. 2016) and Echiura (Lehrke et al. 2012). To our knowledge, Scalibregmatidae stand out as the only annelid taxon so far in which palp-like head appendages appear to have (re-)evolved after an ancestral loss of the structure (see also Martinez et al. 2014).

Nervous system organization in Scalibregmatidae supports this unique evolutionary trajectory as well. As shown for *Travisia* no prominent pro- or peristomial neurite bundles were observed that could be homologized with or presumed as being evolutionary precursors of the either diffuse anterior neural innervation pattern of *S. celticum* or of the distinct set of five palp-like appendage nerves in *A. cavernicola* (see Figs. 3–5). Thus, the lack of such innervating anterior neurite bundles in *Travisia* favors the total loss of palps and the related neuronal scaffold in the sister group of Scalibregmatidae. In *Scalibregma celticum*, nerves arise from almost all regions of the brain (by far the most from the dorsal brain region), including two from the ventral brain region, and extend into the prostomial horns. On the other hand, *A. cavernicola* solely shows five distinct neurite bundles which originate from the dorsal brain region projecting into the palp-like appendages and the prostomial depression. This is comparable to what is observed for antennal innervation but dissimilar to the always both dorsal and ventral innervation of palps in other taxa (Beckers et al. 2019a, b; Kuhl et al. 2022; Kalke et al. 2024).

The prostomial horns of *S. celticum*, which shows a diffuse neuronal scaffold, are not used in feeding (Jumars et al. 2015; Dauer 1983, 1985; Jones 1968; Mortimer & Mackie 2014). Instead, they appear to be putatively sensory, comparable with the “horn-like” appendages in *Tomopteris spp*. (Purschke & Helm 2023). Nevertheless, sensory stout palps in errant annelids are also not directly used for feeding, but contrary to the scalibregmatid prostomial horns, they show a neuronal innervation similar to that of feeding palps (Schmidbaur et al., 2020; Kuhl et al. 2022). In *A. cavernicola*, the innervation pattern - with neurite bundles solely coming from the dorsal root of the circumesophageal connective - resembles that of antennae in e.g., Amphinomidae (Kalke et al. 2024), Eunicida (Kuhl et al. 2022), Nereididae (own observation) and Syllidae (Schmidbaur et al. 2020). In all of these antennal appendages, innervation exclusively originates along the drcc in close proximity to the olfactory nuchal organ nerves, which is true for both scalibregmatid species.

Functional considerations based on our finding about the nervous, but also muscular innervation further support a sensory role of the appendages in Scalibregmatidae. The appendages of the *Axiokebuita–Speleobregma* clade lack a ciliated food-rim typical for feeding-palps and instead generate water currents for indirect food uptake, as previously described by several authors (Martínez et al. 2013; Parapar et al. 2021). The musculature is also poorly developed: only three muscles insert into the appendage or its base, which is in contrast to the complex array of independent palp muscles in nereidids (own observation, data not shown), spirorbids (Rüchel & Müller 2007) or magelonids and spionids (Purschke & Müller 2005). But, although a direct role in feeding comparing to feeding-palps is questionable, these palp-like appendage use the water currents they produce for gathering food particles (own observation), thus indirect feeding. Altogether, these findings suggest that the anterior appendages of *Axiokebuita cavernicola* cannot be homologized with neither grooved or non-grooved feeding-palps nor with sensory palps of other taxa, but evolved convergently as palp-like sensory structures. In *S. celticum* the prostomial horns represent a solely sensory prostomial modification, comparable to the T-shaped prostomium of e.g., *Tomopteris* or some spionids like in the genus *Malacoceros* (Meißner& Götting 2015). Yet, due to the fact that palp-like appendages and prostomial horns in Scalibregmatidae resemble very similar innervation patterns (with a strong neuronal input from the dorsal brain region) both have to be assumed as being homologues structures and an evolutionary morphological transition from prostomial horns to palp-like head appendages seems very likely. Nevertheless, in terms of neuronal and muscular innervation we dismiss the homology of these appendages with the “true” palps of other annelid taxa, contrary to previous statements (Orrhage 1966; Kudenov & Blake 1978; Rouse & Pleijel 2001; Parapar et al. 2011).

However, two ciliated structures in *A. cavernicola* with unclear function or evolutionary history were already mentioned in the literature: the “neck organ” described by Parapar et al. (2011) and the prostomial depression. The “neck organ”, herein interpreted as being a ventral nuchal organ, although ultrastructural data are missing. It is located posterior to the mouth opening and resembles external morphological features like ciliation from paired peristomial pits and the innervation pattern of nuchal organs, which vary greatly across annelids in form and number of ciliated pits. In some cases they appear as slits, or even unpaired so-called caruncles (Purschke 1997; Purschke et al. 2014). Persson & Pleijel (2005) already proposed a sensory function for the “neck organ”/ventral nuchal organ. The chemosensory function of nuchal organs in general was already confirmed by functional analyses in *Platynereis* (Chartier et al. 2018). Our observations show that both, the dorsal and ventral nuchal organ of *A. cavernicola* are innervated by a common nerve that bifurcates before proceeding to each nuchal organ separately. The prominent nerve originates in the dorsal part of the brain, a situation well-known for annelid nuchal organ innervation (Purschke 1997). This finding clearly supports the interpretation of scalibregmatid “neck organ” as being the result of a duplication of the ancestral dorsal nuchal organ, fulfilling the criteria for serial or iterative homology (Schucsich & Wirkner 2007; Wagner 2014).

The second ciliated structure unique to Scalibregmatidae is called “prostomial depression”. The latter is unpaired and located in the center of the prostomium. The dorsal depression is innervated by paired neurite bundles that arise together with the palp-like appendage nerves from the dorsal brain region. A comparable prostomial ciliary tuft was herein observed in *S. celticum*, with a roughly comparable neuronal innervation pattern. However, the highly diffuse and chaotic arrangement of neurites exhibited in *S. celticum* prevents from a clear assignment of the neuronal origin to either the dorsal or ventral roots of the circumoesophageal connectives, and thus a precise homology hypothesis cannot be stated.

In summary, Scalibregmatidae represent a so far unique case in annelid evolution, in which palp-like head appendages appear to have newly evolved after an ancestral loss of palp structures. Neuronal innervation patterns, external appendage morphology, and the muscular scaffold indicate that these structures have to be interpreted as the first example of convergent evolution of annelid palps - an annelid character complex hitherto being regarded as homologous across all annelids. Furthermore, the ventral nuchal organ represents most likely a duplication of the dorsal nuchal organ, whereas the connection between ciliated sensory structures in *S. celticum* and the prostomial depression in *A. cavernicola* remains enigmatic.

## Acknowledgement

The authors want to thank the working group “Animal Evolution and Biodiversity” and “Evolutionary Biology and Ecology” for the financial and technical support.

## Ethics statement

Ethical approval was not required for the study involving animals in accordance with the local legislation and institutional requirements because not applicable by the use of polychaetes.

## Author contributions

PK: Conceptualization, Data curation, Formal analysis, Investigation, Methodology, Visualization, Writing – original draft, review & editing. KW: Data curation, Investigation, Methodology, Resources, Validation, Writing – review & editing. PB: Data curation, Investigation, Methodology, Resources, Validation,– review & editing. AM: review & editing. CH: Conceptualization, Investigation, Project administration, Resources, Supervision, Validation, Writing – review & editing.

## Publisher’s note

All claims expressed in this article are solely those of the authors and do not necessarily represent those of their affiliated organizations, or those of the publisher, the editors and the reviewers. Any product that may be evaluated in this article, or claim that may be made by its manufacturer, is not guaranteed or endorsed by the publisher.

